# Reply to: Commentary on Pang et al. (2023) Nature

**DOI:** 10.1101/2023.10.06.560797

**Authors:** James C. Pang, Kevin M. Aquino, Marianne Oldehinkel, Peter A. Robinson, Ben D. Fulcher, Michael Breakspear, Alex Fornito

## Abstract

In Pang et al. (2023)^1^, we identified a close link between the geometry and function of the human brain by showing that: (1) eigenmodes derived from cortical geometry parsimoniously reconstruct activity patterns recorded with functional magnetic resonance imaging (fMRI); (2) task-evoked cortical activity results from excitations of brain-wide modes with long wavelengths; (3) wave dynamics, constrained by geometry and distance-dependent connectivity, can account for diverse aspects of spontaneous and evoked brain activity; and (4) geometry and function are strongly coupled in the subcortex. Faskowitz et al. (2023)^2^ raise concerns about the framing of our paper and the specificity of the eigenmode reconstructions in result (1). Here, we address these concerns and show how specificity is established by using appropriate benchmarks.

## Main text

Faskowitz et al.’s^2^ critique of our paper’s framing is that it “can be perceived” as a “winner-takes-all… comparison between brain shape and structural connectivity”. This misperception evidently arises from quotes taken out of context and an oversight of the fundamental relationship between geometry and connectivity underlying our approach, as defined formally in Supplementary Information S8 of the original paper^1^. In brief, the use of cortical eigenmodes to model cortical activity rests irrevocably upon a form of distance-dependent connectivity that has been consistently identified in human and non-human data alike^3,4^. In Supplementary Information S1 of this response, we revisit the pertinent details with additional explanatory notes and also clarify how our approach is readily reconciled with lesion studies, as queried by Faskowitz et al.

Faskowitz et al.^2^ also critique the specificity of our geometric eigenmode reconstructions with respect to two proposed criteria. Their first criterion is that geometric eigenmodes “should perform poorly in explaining randomly oriented activity patterns, uncoupled form the underlying cortical anatomy”. To generate such patterns, the authors use the popular spin test^5^, which projects an empirical activation map to a sphere, randomly rotates the map, and then projects the rotated map back onto the cortex (Fig. 1a). The authors show that geometric eigenmodes reconstruct empirical maps of 255 participants from the Human Connectome Project (HCP) and their associated spun versions with comparable accuracy, leading them to conclude that cortical geometry does not constrain function.

**Figure 1.**
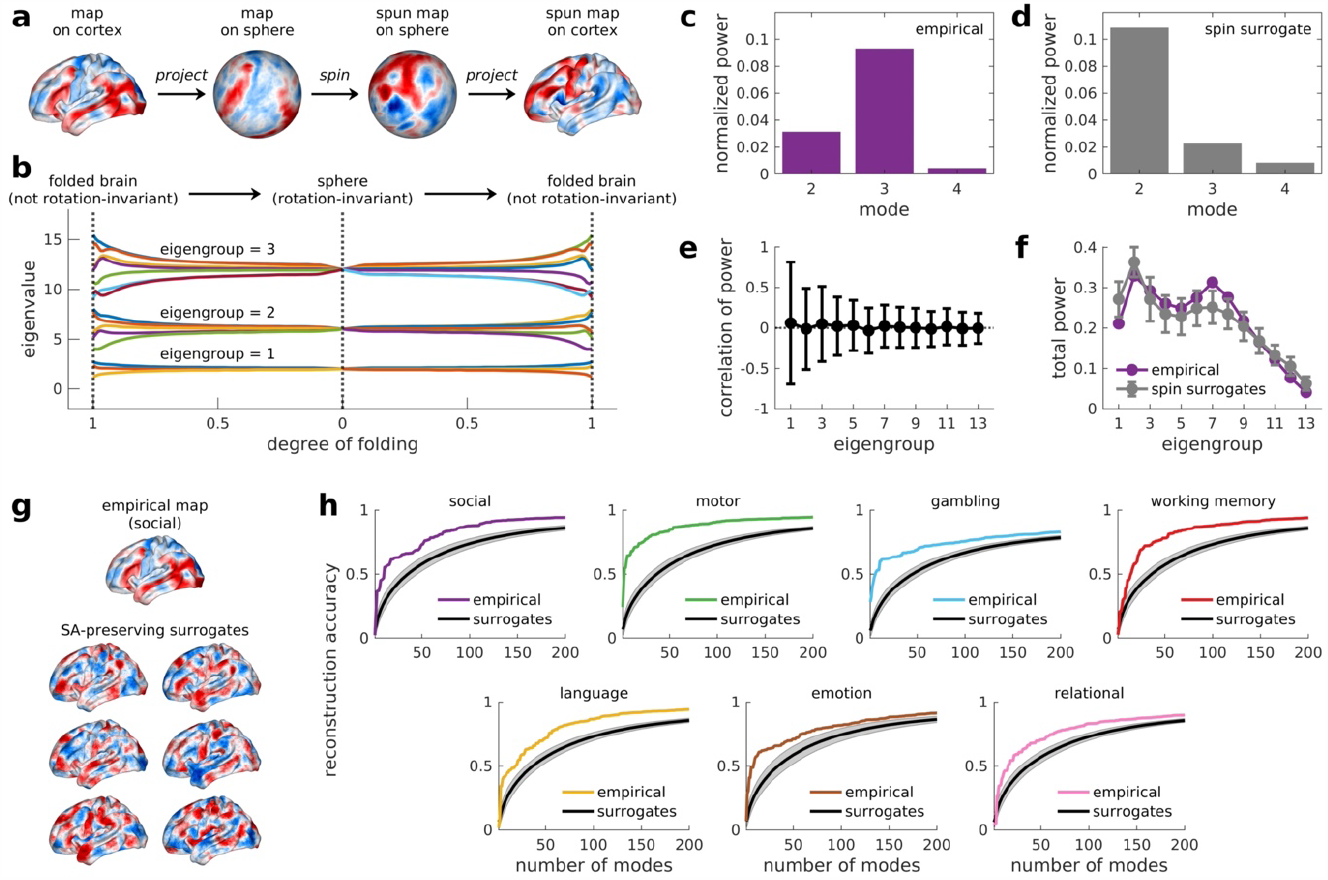
The spin test is inappropriate for inference on geometric reconstructions. **(a)** The spin test projects an empirical activation map to a sphere, spins or rotates the map, and then projects the spun map back onto the cortex. Spinning alters the map’s orientation while preserving its spatial topography. **(b)** The spherical projection distorts cortical geometry, which can be appreciated by observing how the eigenvalues of the cortical geometric eigenmodes change by progressively transitioning from the folded cortex (degree of folding = 1) to a sphere and back to the folded cortex. This panel shows these changes for the first three eigengroups comprising the first 15 non-constant modes, as previously shown in ref. ^7^. In the folded brain, the modes have distinct eigenvalues because the orientations of the modes are “locked in” by cortical geometry, as shown previously^7^ (see also Fig. S3a). As the degree of cortical folding approaches zero (i.e., the cortex becomes a sphere), the eigenvalues within groups converge to their degenerate limit. In this limit, the modes have no preferred orientation and are rotationally-invariant, such that they become interchangeable. Hence, projection back onto the folded brain obtains modes that do not match the original cortical modes (i.e., the colour of each line from top-to-bottom within groups differs from the left to right extremes of cortical folding). **(c**,**d)** Spinning an activation map on a sphere trivially redistributes the power (i.e., coefficient weights of the reconstruction model) across modes within each group. Panel **c** shows the power of modes in the first eigengroup for the empirical map in panel **a**. Panel **d** shows the redistribution of power for the example spun map in panel **a. (e**,**f)** While this redistribution decorrelates the coefficient weights of specific modes for the empirical and spun maps within each group (approximately zero average correlation in panel **e**), the total power of each group, which drives the reconstruction accuracy, is approximately preserved within a comparable wavelength range (panel **f;** see Supplementary Information S2). Hence, the spin test changes the specific coefficient weight of each mode but preserves the overall reconstruction accuracy of the model. Panel **e** shows the mean correlation between mode-specific power within each group, obtained for the empirical map in panel **a** and 1000 surrogate maps. The markers represent the mean and the error bars are the standard deviation. Panel **f** shows the mean total power for each eigengroup, obtained by summing the power within each group and two adjacent groups (to account for leakage between neighbouring groups), for the empirical in panel **a** and 1000 surrogate maps. The markers represent the mean and the error bars are the standard deviation. **(g)** A parametric null model^6^ can be used to generate appropriate surrogate maps that destroy the spatial topography of the empirical map while preserving low-level spatial autocorrelation. **(h)** The accuracy of the geometric eigenmodes in reconstructing the 7 key HCP task-contrast maps is clearly superior to their accuracy in reconstructing the SA-preserving surrogates, thus demonstrating the specificity of the geometric eigenmode model. Panel **b** is adapted from ref. ^7^ with permission.

This conclusion follows from an incorrect application of the spin test, which was developed for inference on pairwise spatial correlations between different maps^5^, not for assessing the relationship of a map with its underlying geometric (or connectomic) support. Projecting data to a sphere distorts cortical geometry, such that the cortical eigenmodes approach spherical harmonics in their degenerate limit (see Fig. 1b). In other words, the specific orientations of the original modes and their distinct eigenvalues, which encode cortical geometry, are absent in the spherical space, rendering the modes interchangeable and rotationally invariant with respect to the activity map. This invariance means that rotating a map in the spherical projection simply redistributes the power (i.e., coefficient weights) of modes within their eigengroups (see Figs. 1c–e and Supplementary Information S2 for details), while approximately conserving the total power of each group (Fig. 1f) and the spatial autocorrelation of the map. Since reconstruction accuracy is determined by the power observed over a given wavelength range, and modes within eigengroups have approximately similar spatial wavelengths, the spin test on the sphere will, by construction, approximately preserve the exact property––reconstruction accuracy––that should be annulled. The spin test is therefore inappropriate for inference on reconstruction accuracy. Note also that the additional null model used by Faskowitz et al.^2^ (Moran spectral randomization) has previously been shown to yield an insufficiently deep randomization of the data^6^ (see Supplementary Information S3 for details).

Proper evaluation of model specificity requires a null benchmark that preserves local, first-order spatial autocorrelations and annuls the spatial topography of the empirical activation map, which captures the influence of geometry on activity. Figures 1g–h show that surrogate maps derived from a parametric null model that satisfies these requirements are reconstructed with lower accuracy than the empirical data (see also Supplementary Information S4 and Figs. S1– S2). Thus, the first specificity criterion proposed by Faskowitz et al.^2^ is fulfilled when appropriate inferential methods are used.

The second specificity criterion proposed by Faskowitz et al.^2^ is that eigenmodes derived from “non-brain-like shapes” should be “less accurate descriptors of cortical activity”. Faskowitz et al.^2^ claim that this criterion is not met because eigenmodes derived from a sphere or a bulbous surface show similar reconstruction accuracy to cortical modes. This analysis simply demonstrates a direct consequence of the coarse-scale geometric similarities between the cortex and a sphere, which have been extensively and formally characterized in prior work^8^. Specifically, ref. ^8^ showed that cortical geometry at large spatial scales (which contains most of the spectral energy of fMRI activation maps; see Fig. 3 in ref. ^1^) can be approximated analytically as a first-order perturbation of spherical geometry, where the perturbations describe the symmetry-breaking effect of cortical folding. As a result, cortical eigenmodes can be expressed as linear combinations of appropriately rotated spherical harmonics^8^ (Figs. 2a–b and Supplementary Information S5). This geometric similarity is why we used spherical approximations to estimate cortical eigenmode wavelengths in our own work (see Eq. (3) in ref. ^1^). In other words, at coarse scales relevant for fMRI, the “non-brain like shapes” used by Faskowitz et al.^2^ are indeed brain-like, and their comparison merely shows that modes obtained from objects with similar geometries reconstruct activity with similar accuracy.

**Figure 2.**
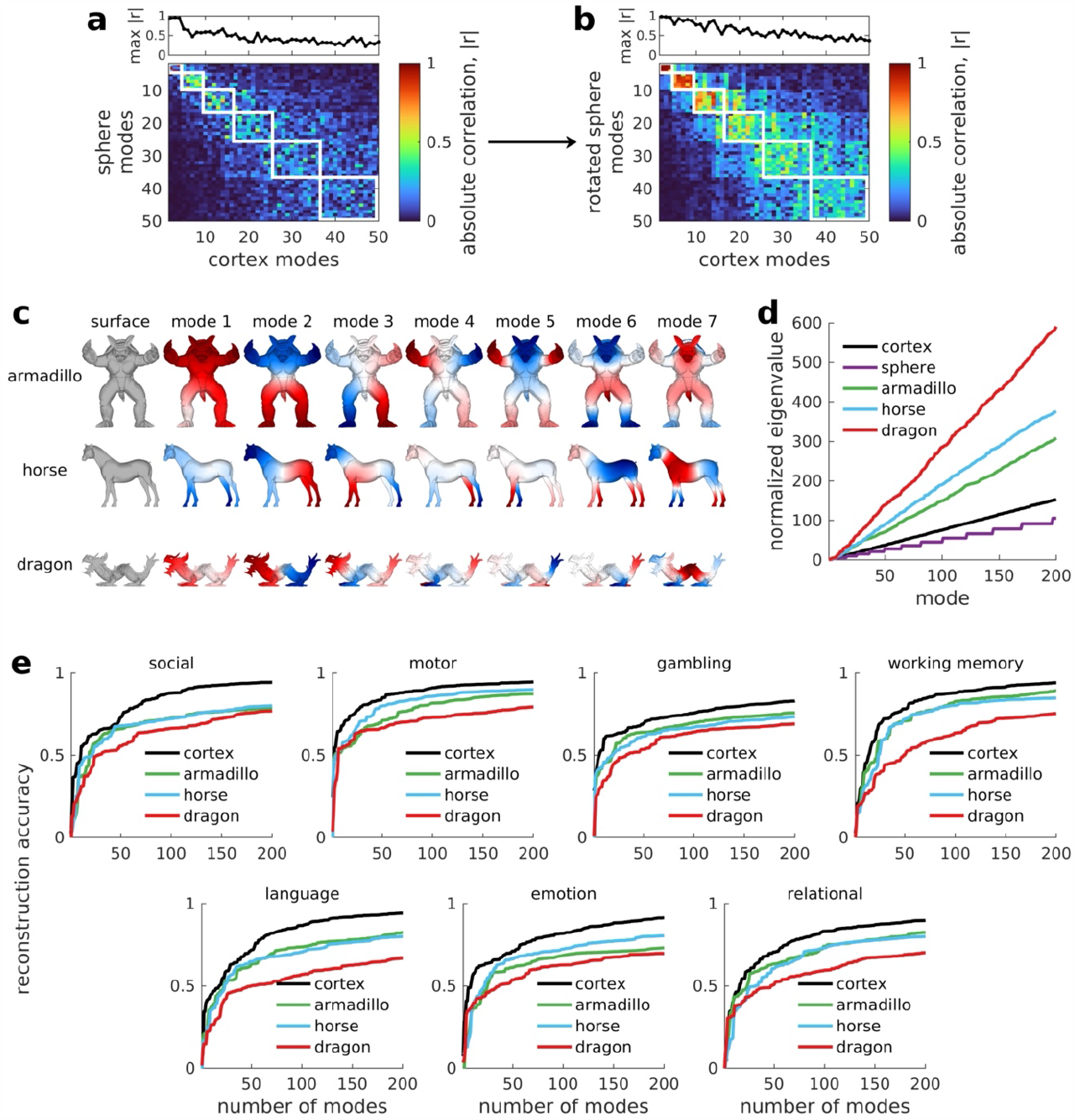
Benchmarking geometric eigenmodes against the eigenmodes of non-brain-like objects. **(a,b)** Absolute spatial correlation, |*r*|, between every pair of cortical and spherical modes before (panel **a**) and after (panel **b**) rotation of the latter. The white boxes represent the eigengroups. The top panels show the highest |*r*| value obtained for each cortical mode, taking into account order flips in each group. The anisotropies of cortical geometry fix the nodal lines of the cortical modes into specific orientations (Fig. S3a). However, spherical modes within each group are degenerate with arbitrary orientations (Figs. 1b and S3b–c) and do not have a one-to-one mapping with the cortical modes, resulting in lower correlations in panel **a**. After rotating the spherical modes in panel **b**, the correlations with cortical modes are higher for the first 6 groups, which account for ∼70% of the reconstruction accuracy of geometric eigenmodes (see Fig. 1d of ref. ^1^). Thus, as previously shown^8^, low-order cortical modes can be approximated in terms of spherical modes, allowing them to have comparable reconstruction accuracies and explaining the effect shown by Faskowitz et al.^2^. **(c)** We more appropriately benchmark the specificity of the geometric eigenmodes against the eigenmodes of three non-brain-like objects (see Supplementary Information S5 for details). This panel shows the surface meshes of these objects (armadillo, horse, and dragon) and their eigenmodes. **(d)** Normalized eigenvalue spectra of the three objects, cortex, and sphere. Note that more similar objects will have more similar spectra. The cortex and spherical modes have more similar spectra compared to the other objects. Moreover, the spectra of the three non-brain objects increase more steeply due to their jagged protrusions (e.g., legs, tails, spikes), which cause high-frequency geometric fluctuations (see also Fig. S4). **(e)** The accuracy of the cortical geometric eigenmodes in reconstructing the 7 key HCP task-contrast maps is higher than those of the 3 non-brain objects, thus demonstrating the specificity of the geometric eigenmode model.

In Figs. 2c–e, we show that eigenmodes derived from objects with clearly “non-brain-like” coarse-scale geometries (i.e., shapes with ridges, sharp peaks, and asymmetries) yield poorer reconstruction accuracies than the eigenmodes of the cortex (see Supplementary Information S5 for details). We thus fulfill Faskowitz et al.’s^2^ second specificity criterion when using appropriate benchmarks.

We further emphasize that the similar performance of sphere-like and cortical modes does not imply that cortical geometry is unrelated to function. A drumhead and a pancake have similar geometries and one could plausibly reconstruct the vibrational patterns of the drum using the geometric modes of the pancake. Nonetheless, the drumhead’s vibrations are still constrained by its geometry––the geometric features of a system will influence its dynamics even if those features are shared with other objects. For this reason, comparisons with arbitrary geometric models offer limited insights. As stated in ref. ^1^, our goal was to identify a parsimonious basis set of physically constrained eigenmodes, not an arbitrary basis set that is optimal in some purely statistical sense. Alternative phenomenological (i.e., non-physiological) basis sets may reconstruct fMRI data at equivalent accuracy but they do not shed light on the anatomical constraints or generative processes that give rise to brain activity^9^. Geometric eigenmodes are derived from a physical property of the brain and are formally related to dynamics through neural field theory^10–12^, which has explained a diverse array of neurophysiological findings over several decades (see for example refs. ^13–15^). Such physically principled basis sets should always be prioritized because they offer privileged insights into generative mechanisms. Their specificity is therefore optimally established with respect to other physiologically-plausible basis sets, as already demonstrated in our original analysis^1^.

In summary, in Pang et al. (2023)^1^, we used multiple lines of converging evidence to show that cortical geometry, distance-dependent connectivity, and wave dynamics are sufficient to explain diverse patterns of brain activity evident in electrophysiological and fMRI data^1^. Faskowitz et al.^2^ question one line of evidence, asking whether eigenmode-based reconstructions show sufficient specificity. Here, we show that the authors’ proposed specificity criteria are met when appropriate benchmarks are used. We further caution against interpretations of our work based on a simplistic dichotomy between geometry and connectivity. The relevance of geometry for function comes from an explicit biophysical model of neuronal dynamics, which assumes a specific form of distance-dependent connectivity that dominates empirical connectome data^3,4^ (see Supplementary Information S1). While this exponential distance-dependence effect does not capture all aspects of brain connectivity, our findings^1^ indicate that it is sufficient to parsimoniously explain diverse dynamical phenomena measured with classical fMRI paradigms.

## Data availability

Raw and pre-processed HCP data can be accessed at https://db.humanconnectome.org/. All source data to generate the results of the manuscript are openly available at https://github.com/NSBLab/BrainEigenmodes and https://osf.io/xczmp/.

## Code availability

Computer codes to analyse results and reproduce the figures of the manuscript are openly available at https://github.com/NSBLab/BrainEigenmodes.

## Acknowledgements

HCP data were provided by the Human Connectome Project, Wu-Minn Consortium (Principal Investigators: David Van Essen and Kamil Ugurbil; 1U54MH091657) funded by the 16 NIH Institutes and Centers that support the NIH Blueprint for Neuroscience Research, and by the McDonnell Center for Systems Neuroscience at Washington University. This work was supported by the MASSIVE HPC facility (www.massive.og.au), Sylvia and Charles Viertel Foundation grant 2017042 to A.F., National Health and Medical Research Council grants 1197431 and 1146292 to A.F., Australian Research Council grant DP200103509 to A.F., Australian Research Council Laureate Fellowship FL220100184 to A.F., National Health and Medical Research Council grant 2008612 to M.B., Australian Research Council Laureate Fellowship FL140100025 to P.A.R., and Australian Research Council Center of Excellence CE140100007 to P.A.R.

## Author contributions

J.C.P., M.B., and A.F. designed the methodology. J.C.P. and A.F. performed the investigation and administered the project. J.C.P. developed the visualizations. A.F. acquired funding and supervised the project. J.C.P. and A.F. wrote the original draft. All authors reviewed and edited the final manuscript.

## References

1. Pang, J. C. et al. Geometric constraints on human brain function. Nature 618, 566–574 (2023).

2. Faskowitz, J. et al. Commentary on Pang et al. (2023) Nature. 2023.07.20.549785 Preprint at 10.1101/2023.07.20.549785 (2023).

3. Markov, N. T. et al. Cortical High-Density Counterstream Architectures. Science 342, 1238406 (2013).

4. Roberts, J. A. et al. The contribution of geometry to the human connectome. NeuroImage 124, 379–393 (2016).

5. Alexander-Bloch, A. F. et al. On testing for spatial correspondence between maps of human brain structure and function. NeuroImage 178, 540–551 (2018).

6. Burt, J. B., Helmer, M., Shinn, M., Anticevic, A. & Murray, J. D. Generative modeling of brain maps with spatial autocorrelation. NeuroImage 220, 117038 (2020).

7. Robinson, P. A. et al. Eigenmodes of brain activity: Neural field theory predictions and comparison with experiment. NeuroImage 142, 79–98 (2016).

8. Gabay, N. C. & Robinson, P. A. Cortical geometry as a determinant of brain activity eigenmodes: Neural field analysis. Physical Review E 96, (2017).

9. Shinn, M. Phantom oscillations in principal component analysis. 2023.06.20.545619 Preprint at 10.1101/2023.06.20.545619 (2023).

10. Wright, J. J. & Liley, D. T. J. Simulation of electrocortical waves. Biological Cybernetics 72, 347–356 (1995).

11. Jirsa, V. & Haken, H. Field Theory of Electromagnetic Brain Activity. Physical Review Letters 77, 960–963 (1996).

12. Robinson, P. A., Rennie, C. J. & Wright, J. J. Propagation and stability of waves of electrical activity in the cerebral cortex. Physical Review E 56, 826–840 (1997).

13. Robinson, P. A. et al. Prediction of electroencephalographic spectra from neurophysiology. Physical Review E 63, 021903 (2001).

14. Abeysuriya, R. G., Rennie, C. J. & Robinson, P. A. Physiologically based arousal state estimation and dynamics. Journal of Neuroscience Methods 253, 55–69 (2015).

15. Gabay, N. C., Babaie-Janvier, T. & Robinson, P. A. Dynamics of cortical activity eigenmodes including standing, traveling, and rotating waves. Phys. Rev. E 98, 042413 (2018).

