## Supplementary Information for "Reply to: Commentary on Pang et al. (2023) Nature"

### S1. The relationship between geometry, connectivity, and dynamics

Faskowitz et al.[1] state that the central claim of our work in ref. [2] is that "the geometry of the cortical surface provides a more parsimonious explanation of brain activity than structural brain connectivity" and that this claim "rests largely on a comparison between brain shape and connectivity" that can be perceived as "winner-takes-all". This characterization of our work is inaccurate and appears to follow from quotes taken out of context and an oversight of the formal links between geometry and connectivity established in Supplementary Information S8 of our original paper[2]. In this section, we rehearse and further explain the formal relationship between geometry and connectivity that underpins our approach.

The constraining effect of geometry on spatiotemporal brain activity derives from an established class of biophysical models called neural field theory (NFT)[3–5]. In NFT, brain dynamics on mesoscopic and macroscopic spatial scales (i.e., >0.5 mm) are treated as local averages of the activity of neural populations (e.g., firing rate). A physiologically constrained

NFT developed by Robinson and colleagues[5] further treats the resulting mean-field activity as traveling waves of excitation that propagate through a continuous medium[5–7]. This continuum approximation is possible because of the dense cellular packing and horizontal connectivity of the cortex[8]. Hence, continuous wave-like dynamics arise at sufficiently coarse spatial scales and have been widely observed in local field potentials[9–12], electro- and magneto-encephalography[13,14], and functional magnetic resonance imaging (fMRI)[15–17].

In the Robinson et al. NFT model, neural activity $\phi(\boldsymbol{r},t)$ at location $\boldsymbol{r}$ and time $t$ is described by the isotropic damped wave equation without regeneration[4,5,18]

$$\left[\frac{1}{\gamma_s^2}\frac{\partial^2}{\partial t^2}+\frac{2}{\gamma_s}\frac{\partial}{\partial t}+1-r_s^2\nabla^2\right]\phi(\boldsymbol{r},t)=Q(\boldsymbol{r},t), \qquad (S1)$$

where $Q$ is an external input, $\gamma_s$ is the damping rate, and $r_s$ is spatial length scale wave propagation. Equation (S1) can be solved using the Green's function approach such that

$$\phi(\boldsymbol{r},t)=\int W(\boldsymbol{r},t;\boldsymbol{r}',t')Q(\boldsymbol{r}',t')d^2\boldsymbol{r}'dt', \qquad (S2)$$

where the kernel $W(\boldsymbol{r},t;\boldsymbol{r}',t')$, also known as the propagator or Green's function, defines how activity between points and time influence each other, specifically via the white matter connectivity. Mathematically, $W$ is the solution of Eq. (S1) with delta-function stimulus $Q(\boldsymbol{r},t)=\delta(\boldsymbol{r})\delta(t)$, that is the spatiotemporal kernel to get the impulse response.

Eq. (S2) states that the activity, $\phi$, at location $\boldsymbol{r}$ and time $t$ is a convolution of the external input, $Q$, and connectivity with other areas, as defined by the kernel, $W$. The kernel $W$ can take a variety of forms[5,19]. In our work[2], we assume an isotropic case such that $W$only depends on the spatial separation between points and the time difference such that $W(\boldsymbol{r},t;\boldsymbol{r}',t')=W(\boldsymbol{r}-\boldsymbol{r}',t-t')$, and decays exponentially as a function of physical distance. We use this kernel following (1) extensive evidence for an exponential distance rule (EDR) in white matter connectivity across multiple modalities (tractography, tracer-based studies) and species (humans, other primates, rodents)[20–23]; and (2) extensive prior work showing that the kernel is effective in explaining numerous empirical phenomena observed in electroencephalography (EEG) recordings[24–26]. We thus seek to evaluate the extent to which such a simple approximation can explain diverse fMRI findings. To explicitly show the form of $W$ we use, we will briefly show its derivation based on ref. [5,27].

We first take the Fourier transform of Eq. (S1) with $\phi(\boldsymbol{r},t)=W(\boldsymbol{r}-\boldsymbol{r}',t-t')$ and $Q(\boldsymbol{r},t)=\delta(\boldsymbol{r})\delta(t)$ such that,

$$\left[-\frac{\omega^2}{\gamma_s^2}-\frac{2i\omega}{\gamma_s}+1+r_s^2k^2\right]W(\boldsymbol{k},\omega)=1, \qquad (S3)$$

where $\omega$ is the temporal angular frequency, $k=|\boldsymbol{k}|$ is the spatial wavenumber, and $W(\boldsymbol{k},\omega)$ is the Fourier transform of $W(\boldsymbol{r}-\boldsymbol{r}',t-t')$. Thus,

$$W(\boldsymbol{k},\omega)=\frac{1}{-\frac{\omega^2}{\gamma_s^2}-\frac{2i\omega}{\gamma_s}+1+r_s^2k^2}, \qquad (S4)$$

$$W(\boldsymbol{k},\omega) = \frac{1}{(1 - i\omega/\gamma_S)^2 + r_s^2 k^2}. \qquad (S5)$$

Taking the inverse Fourier transform of Eq. (S5) gives,

$$W(\boldsymbol{r} - \boldsymbol{r}', t - t') = \iint \frac{e^{i\boldsymbol{k}\cdot(r - r') - i\omega(t - t')}}{(1 - i\omega/\gamma_S)^2 + r_s^2 k^2} \frac{\mathrm{d}^2\boldsymbol{k}}{(2\pi)^2} \frac{\mathrm{d}\omega}{2\pi}, \qquad (S6)$$

$$W(\boldsymbol{r} - \boldsymbol{r}', t - t') = \int_0^\infty \frac{\mathrm{d}k}{2\pi} k J_0(k|\boldsymbol{r} - \boldsymbol{r}'|) \int \frac{\mathrm{d}\omega}{2\pi} \frac{e^{-i\omega(t - t')}}{(1 - i\omega/\gamma_S)^2 + r_s^2 k^2}, \qquad (S7)$$

where $J_0$ is the zeroth-order Bessel function of the first kind. After further algebraic manipulations, Eq. (S7) can be further simplified giving,

$$W(\boldsymbol{r} - \boldsymbol{r}', t - t') = \frac{\gamma_s}{r_s} \frac{\exp\left(\frac{-|\boldsymbol{r} - \boldsymbol{r}'|}{r_s}\right)}{\sqrt{\gamma_s^2 r_s^2 (t - t')^2 - |\boldsymbol{r} - \boldsymbol{r}'|^2}} \Theta[\gamma_s r_s (t - t') - |\boldsymbol{r} - \boldsymbol{r}'|], \qquad (S8)$$

where $\Theta$ is the Heaviside step function. Equation (S8) indicates that the connectivity between two cortical locations, $\boldsymbol{r}$ and $\boldsymbol{r}'$, decays approximately exponentially as a function of their distance, $|\boldsymbol{r} - \boldsymbol{r}'|$, based on their isotropic anatomical connectivity. More formally, the distance-dependence corresponds to the zeroth-order modified Bessel function of the second kind, $K_0(|\boldsymbol{r} - \boldsymbol{r}'|/r_s)$, which can be obtained by integrating $W(\boldsymbol{r} - \boldsymbol{r}', t - t')$ with respect to the time difference $t - t'$; see ref. [5] for further details.

Substituting Eq. (S8) into Eq. (S2) allows us to calculate the activity at location $\boldsymbol{r}$ and time $t$ in integral form as,

$$\phi(\boldsymbol{r}, t) = \frac{\gamma_s}{r_s} \int \frac{\exp\left(\frac{-|\boldsymbol{r} - \boldsymbol{r}'|}{r_s}\right)}{\sqrt{\gamma_s^2 r_s^2 (t - t')^2 - |\boldsymbol{r} - \boldsymbol{r}'|^2}} \Theta[\gamma_s r_s (t - t') - |\boldsymbol{r} - \boldsymbol{r}'|] Q(\boldsymbol{r}', t') d^2\boldsymbol{r}' dt', \qquad (S9)$$

which shows the explicit contribution of connectivity. Note that the solution of Eq. (S1) is the same as the solution of Eq. (S9), but the latter form shows the connectivity inherent in our geometric model, which is not readily seen in the former expression.

This derivation thus shows explicitly how the wave dynamics given by Eq. (S1) depend on EDR-like connectivity. We refer interested readers to ref. [5] for detailed derivations and note that Eqs. (S1), (S2), (S8), and (S9) respectively correspond to Eqs. (S6), (S7), (S8), and (S9) in the original Supplementary Information of our paper[2]. Note that choosing a different form of $W$ for Eq. (S2) yields a different partial differential equation (PDE) to Eq. (S1). For example, a purely local square footprint (i.e., purely local nearest-neighbor connectivity) of the type Faskowitz et al. imply that we assume, yields a very different PDE. Techniques that are appropriate for studying Eq. (S1), such as traveling wave solutions, are no longer appropriate[28].

We next rehearse how the geometric eigenmodes are linked to the NFT-based neural activity $\phi(\boldsymbol{r}, t)$ and connectivity. A useful way to solve the wave equation in Eq. (S1) is via the separation of variables approach used in calculus, which applies to systems with spatial properties that do not change at time scales relevant to the system's short-term dynamics. Hence, we can make the ansatz that $\phi(\boldsymbol{r},t)$ can be decomposed into functions of space and time separately such that,

$$\phi(\boldsymbol{r},t) = \phi(\boldsymbol{r})\phi(t). \qquad (S10)$$

We then substitute Eq. (S10) into Eq. (S1). For simplicity, we only show here the steps for the case of $Q(\boldsymbol{r},t) = 0$, but the mathematics can be extended for general cases. Hence,

$$\left[\frac{1}{\gamma_s^2}\frac{\partial^2}{\partial t^2} + \frac{2}{\gamma_s}\frac{\partial}{\partial t} + 1 - r_s^2\nabla^2\right]\phi(\boldsymbol{r})\phi(t) = 0, \qquad (S11)$$

$$\phi(\boldsymbol{r})\left[\frac{1}{\gamma_s^2}\frac{d^2\phi(t)}{dt^2} + \frac{2}{\gamma_s}\frac{d\phi(t)}{dt} + \phi(t)\right] = r_s^2\phi(t)\nabla^2\phi(\boldsymbol{r}). \qquad (S12)$$

Note that the temporal partial derivatives in Eq. (S11) become ordinary derivatives in Eq. (S12) because $\phi(t)$ is a function of $t$ only. Dividing Eq. (S12) by $r_s^2\phi(\boldsymbol{r})\phi(t)$ yields,

$$\frac{1}{r_s^2\phi(t)}\left[\frac{1}{\gamma_s^2}\frac{d^2\phi(t)}{dt^2} + \frac{2}{\gamma_s}\frac{d\phi(t)}{dt} + \phi(t)\right] = \frac{\nabla^2\phi(\boldsymbol{r})}{\phi(\boldsymbol{r})}. \qquad (S13)$$

Because the left- and right-hand sides of Eq. (S13) are independent of $\boldsymbol{r}$ and $t$, respectively, the only permittable solutions are when both sides are equal to a common constant. We assume the constant to be $-\lambda$, where $\lambda > 0$, to avoid diverging solutions. Therefore, Eq. (S13) separates into two equations,

$$\frac{1}{\gamma_s^2}\frac{d^2\phi(t)}{dt^2} + \frac{2}{\gamma_s}\frac{d\phi(t)}{dt} + (1 + r_s^2\lambda)\phi(t) = 0, \qquad (S14)$$

$$\nabla^2\phi(\boldsymbol{r}) = -\lambda\phi(\boldsymbol{r}). \qquad (S15)$$

Note that $\nabla$ is the Laplace-Beltrami operator (LBO) and Eq. (S15) is the well-known Helmholtz equation used to calculate the geometric eigenmodes in our work. The functions $\phi(\boldsymbol{r})$ correspond to the geometric eigenmodes, which correspond to the spatial component of the solutions to the NFT wave equation. Moreover, because $\phi$ depends on the EDR-like connectivity kernel $W$ defined in Eq. (S4), there is an intimate relationship between the geometric eigenmodes and connectivity and thus between geometry and connectivity; they are two sides of the same coin. For these reasons, Faskowitz et al.'s[1] framing of our analysis as "a comparison between brain shape and connectivity" that "can be perceived as winner-takes-all" is inaccurate. Our geometric model incorporates a specific form of connectivity, approximated as an isotropic, EDR-like connectivity. Indeed, this close relationship between geometry and connectivity is precisely why we evaluated eigenmodes extracted from networks derived from a stochastic, EDR-like rule (see ref. [2], p. 568).

While we agree with Faskowitz et al.[1] that "a simple proximity rule fails to account for the marked heterogeneity and specificity of macroscale white matter connectivity", an EDR-like process can nonetheless explain many aspects of connectome architecture [20–23,29–31] and thus represents a suitable first-order approximation. A key finding of our analysis in Pang et al.[2] is that this approximation is sufficient to explain diverse neurophysiological phenomena. We further emphasize that classical discrete, connectome-informed approaches also make approximations. The two approaches simply differ in the specific approximations that they make and the anatomical properties that they prioritize. Geometric eigenmodes prioritize the dense local, horizontal connectivity of the cortex and physical constraints (distance, geometry) on activity, but rely on a simple approximation of inter-regional connectivity. Connectome eigenmodes prioritize heterogeneous and topologically complex inter-regional connectivity, but do not directly account for local, horizontal connectivity or physical constraints. The superiority of the geometric eigenmodes and the wave model in our original analysis highlight the importance of local, horizontal connections and physical constraints in shaping patterned dynamics. As we stated in ref. [2], these results do not mean that specific, heterogeneous, and topologically complex connections (i.e., connections beyond EDR) are unimportant for brain function. Rather, our findings indicate that classical fMRI paradigms do not readily reveal the dynamical effects of such connections when assessed with eigenmode expansions or biophysical models. This is perhaps unsurprising given the low spatiotemporal resolution of fMRI and estimates that ~80% of synaptic inputs within a cortical area correspond to recurrent, intra-regional connections that predominantly originate from within a 2 mm radius[32]—precisely those connections that are not captured well by discrete macroscopic connectome models. These points were acknowledged on p. 572 of ref. [2].

Having established a formal link between geometry and connectivity, we can now address Faskowitz et al.'s[1] mandate that the effects of geometry "must be reconciled with a century's worth of observations wherein direct insults to white matter pathways leave surface geometry intact but nonetheless result in acute changes in function, behaviour, and cognition". It should be clear from the preceding discussion that this comment rests on a false dichotomy between geometry and connectivity. It also overlooks formal theory already developed to address this very topic [33]. Specifically, lesions will perturb the connectivity kernel, $W$, causing spatial heterogeneity and anisotropies of regional connectivity. As a result, the spatial structure of the geometric eigenmodes and their corresponding eigenvalues will change to account for this heterogeneity[18,33], which ultimately changes the resulting dynamics. Ref. [33] has theoretically shown that the effect of acute lesions and other modifications (e.g., due to drugs or injury) can be approximated as a first-order perturbation on the connectivity kernel such that $W = W^{(0)} + W^{(1)}$, where $W^{(0)}$ is the original, unperturbed kernel defined in Eq. (S8) and $W^{(1)}$ is the perturbed kernel capturing the brain modifications. As an example, white-matter lesions will take forms of $W^{(1)}$ that capture changes in the strength of connections between distinct pairs of locations $\boldsymbol{r}$ and $\boldsymbol{r}'$, whilst grey-matter lesions perturb the surface geometry and thus change the strength of connections between location $\boldsymbol{r}$ (the lesion location) and all $\boldsymbol{r}'$. Both lesion types will change the overall structure of $W$, directly impacting the activity field $\phi$ and the geometric constraints on dynamics, resulting in acute changes in function, cognition, and behaviour. We look forward to future work building on this approach and other theoretical developments[31] to investigate such scenarios.

### S2. Limitations of the spin test

The spin test[34] was originally developed for statistical inference on spatial correlations between two cortical maps. The intrinsic spatial autocorrelation of MRI data can inflate spatial correlations and violates the independence assumptions of traditional parametric inferential tests. The spin test provides a null model that breaks the spatial dependence between the two maps by rotating or spinning one map while preserving its topography (and thus, its spatial autocorrelation). However, the geometric anisotropy of the cortex makes it difficult to rotate maps on the cortical surface, so they are first projected to a sphere. The rotation is applied in spherical space before the rotated map is projected back onto the cortex. This procedure offers a valid method for generating a null ensemble for inference on pairwise correlations between spatial maps. In this case, the spin test appropriately annuls the object of inference—the spatial correlation between two maps—while preserving other low-level features of the data, such as their spatial autocorrelation.

Faskowitz et al.[1] used the spin test for inference on the accuracy of the geometric eigenmode expansion in reconstructing Human Connectome Project (HCP) data[35,36]. It is important to note that the object of inference here is no longer the spatial correlation between two maps but the accuracy of a model that reconstructs an empirical map from a weighted sum of eigenmodes. This distinction creates a problem when the rotation is applied in spherical space because, by construction, the spherical projection distorts the cortical geometry so that it becomes sphere-like. The resulting geometric eigenmodes of the cortex are also distorted, such that they approximate the harmonics of the sphere. This effect can be appreciated by considering the first three non-global eigenmodes of the cortex and a sphere (Figs. S3a–c). In the cortex, these modes approximately represent spatial variations along the rostro-caudal, dorso-ventral, and medio-lateral directions (Fig. S3a). In the sphere, one can orient these modes along the same axes, although this alignment is arbitrary because the sphere is symmetric such that the rostral, caudal, dorsal, ventral, medial, and lateral poles are undefined (this is why spherical modes projected on the cortex do not match the real cortical modes; Fig. S3c) Thus, spherical eigenmodes are rotationally invariant with degenerate (i.e., equivalent) eigenvalues (Fig. 1b). This degeneracy allows one to separate the spherical modes into harmonic groups, or “eigengroups”, describing spatial variations of the same wavelength. Eigenmodes are interchangeable within these groups.

As shown previously[37], anisotropies of cortical geometry “lock in” the nodal lines of the modes, thereby fixing their orientation (Fig. S3a). This is clearly evident as one tracks the eigenvalues of different eigengroups as the cortex is gradually deformed to a sphere and back again (Fig. 1b). Figure 1b was adapted with permission from ref. [38], which was obtained by plotting the eigenvalue solutions of Eq. (S15) starting from the folded cortex up to a sphere and back to the folded cortex. The degree of folding (x-axis of Fig. 1b) represents the fraction of the inflation algorithm steps remaining in going from the cortex to a sphere. In the cortex, modes within the same group have different eigenvalues but they gradually converge to their rotationally-invariant, degenerate limit when the cortical folds are completely removed (degree of folding = 0); i.e., in the final spherical projection. This convergence means that the activation map also becomes rotationally invariant. Thus, any rotation of the activation map will always be reconstructed with the same accuracy because the map’s spatial topography and its inherent spatial wavelength are unchanged. The rotation only redistributes the energy or power (i.e., coefficient weights) across different modes (Figs. 1c–d).

Figure 1c shows the normalized power for the empirical social task map of the HCP data and Fig. 1d shows one example of its corresponding spin test surrogate. The distribution of power across the groups is clearly different, as demonstrated in Fig. 1e, which shows the correlation of the power distribution in each group for the empirical map in Fig. 1a and 1000 surrogate maps. However, the total power within a given group (Fig. 1f) is approximately conserved, with some leakage to adjacent groups, which we have taken into account in Fig. 1f by taking the sum of the power in a group and its two adjacent groups (note that eigengroup 1 only has one adjacent group; i.e., eigengroup 2). The reconstruction accuracy of the eigenmode expansion at a particular mode order is determined by the power of the modes spanning the corresponding wavelength range, such that the spin test will, by construction, approximately preserve the exact property that must be annulled for sensible inference (i.e., model reconstruction accuracy). For this reason, the spin test should not be used for inference on model reconstruction accuracy. The spin test is a suitable method for inference on the coefficient weights themselves, but this form of inference was not performed in our original analysis[2].

**S3. Limitations of the Moran spectral randomization model**

Faskowitz et al.[1] employed a second null model (i.e., Moran spectral randomization[39–41]) to support their claims based on the spin test. Moran spectral randomization involves constructing a spatial weight matrix defining relationships between points directly on the cortex (typically based on distances), thus avoiding spherical projection. The weight matrix is decomposed to obtain a basis set of orthogonal spatial eigenvectors called Moran eigenvectors. The target map (in this case, the HCP empirical activation maps) are projected onto the Moran eigenvectors, and the signs of the projection weights are randomized to produce a null surrogate for the target map. However, when the target map loads strongly onto one or more of the Moran eigenvectors, prior work[40] has shown that this approach produces surrogate maps that carry the dominant spatial features of the empirical map regardless of the random flipping of the signs of the weights. Therefore, the surrogates obtained from this null model can be strongly correlated or anticorrelated with the target map (see Supplementary Fig. 7 in ref. [40]), meaning that the approach does not generate a sufficiently deep randomization of the data. Moran spectral randomization is thus inappropriate for evaluating the efficacy of the geometric eigenmode reconstruction.

**S4. BrainSMASH – a parametric null model for spatial inference**

An appropriate inferential procedure for testing geometric specify requires a null model that disrupts the spatial topography of the empirical map to yield a pattern "uncoupled form the underlying cortical anatomy"[1]. This is precisely the null model offered by the method developed in the Brain Surrogate Maps with Autocorrelated Spatial Heterogeneity (BrainSMASH) platform[40]. The method produces surrogate maps with spatial autocorrelation (SA) matched to the SA of the target map. Briefly, the approach first calculates the variogram of the target map, which quantifies the variance between all pairwise points as a function of distance. For example, a map comprising pure white noise will have a flat variogram. The target map is then randomly shuffled, effectively destroying its spatial structure and SA. SA is reintroduced by smoothing the randomly shuffled map with a distance-dependent kernel (e.g., exponential). The variogram of the smoothed random map is calculated, regressed onto the target map's variogram, and the variogram fit is assessed. This process is repeated until the best-fitting variogram is found.

We applied the above method to create 1000 SA-preserving surrogate maps for each of the 7 key HCP task-contrast maps. Examples are shown in Fig. 1g. The resulting surrogates are properly randomized, with narrow, zero-centered distributions of correlations when compared to the empirical maps (Fig. S1a), but with a good fit to the SA of the empirical data (Fig. S1b). Moreover, the surrogates also appropriately annul the spatial wavelength structure of the empirical data, with the power spectra of the modes approximately whitened (Fig. S1c), appropriately destroying the dominant long-wavelength characteristics we previously found for most fMRI task-activation maps (see Fig. 3 in ref. [2]).

Comparisons of reconstruction accuracies obtained for the empirical maps and the surrogate maps generated using this parametric null model are presented in Fig. 1h. Following Faskowitz et al.[1], we also performed statistical inference by calculating the *p*-values of the reconstruction accuracy, which represent the proportion of times the accuracy of the geometric eigenmodes in reconstructing the empirical maps exceeds the accuracy in reconstructing the surrogate maps (Fig. S2). The results clearly show that the geometric eigenmode model reconstructs real empirical data with greater accuracy than the surrogate maps.

**S5. The specificity of cortical geometry with respect to other shapes**

Prior work has extensively characterized geometric similarities between the cortex and sphere[37,38]. Specifically, ref. [37] used first-order perturbation theory to derive an analytic approximation of non-constant cortical modes $\left|y_{\lambda\mu}^{(1)}\right\rangle$ as a linear combination of appropriately rotated spherical harmonics $|Y_l^m\rangle$ of order $m$ and degree $l$, given by

$$\left|y_{\lambda\mu}^{(1)}\right\rangle = \sum_{l\neq\lambda}\sum_{m}\frac{\left\langle Y_l^m\middle|\nabla_p^2\middle|y_{\lambda\mu}^{(0)}\right\rangle}{E_{\lambda\mu}^{(0)}-E_{lm}}\left[|Y_l^m\rangle+\sum_{j\neq\mu}\frac{\left\langle y_{\lambda j}^{(0)}\middle|\nabla_p^2\middle|Y_l^m\right\rangle}{E_{\lambda j}^{(1)}-E_{\lambda\mu}^{(1)}}\left|y_{\lambda j}^{(0)}\right\rangle\right], \qquad (S16)$$

where $\left|y_{\lambda\mu}^{(1)}\right\rangle$ and $\left|y_{\lambda\mu}^{(0)}\right\rangle$ are the first-order perturbed and unperturbed eigenmodes of degree $\lambda$ and order $\mu$, respectively, written in Dirac bra-ket notation. The former corresponds to the cortical eigenmodes and the latter are rotated versions of spherical harmonics of degree $\lambda$. Note that the indices $\lambda$ and $\mu$ are used to match the $l$ and $m$ indices for the spherical harmonics, with the degree, $\lambda$, corresponding to the eigengroup discussed in Section S2 and $\mu$ being the index of the eigenmodes comprising the eigengroup. The term $\nabla_p^2$ is the Laplace-Beltrami operator for the small perturbation due to cortical folding and $E_{\lambda\mu}^{(1)}$, $E_{\lambda\mu}^{(0)}$, and $E_{lm}$ correspond to the perturbed, unperturbed, and spherical harmonic eigenvalues, respectively.

Equation (S16) shows that cortical eigenmodes are a linear combination of spherical harmonics (first term of the right-hand side) and unperturbed eigenmodes (second term of the right-hand side), the latter of which are just rotated versions of the spherical harmonics. It is also important to highlight that the contribution of the spherical harmonics to the perturbed eigenmodes is scaled by the inverse of $E_{\lambda\mu}^{(0)} - E_{lm}$; hence, the contribution of increasingly higher-order spherical harmonics (harmonics with higher eigenvalues and that vary over smaller wavelengths) becomes smaller and lower-order spherical harmonics dominate. We refer interested readers to ref. [37] for detailed derivations and discussion.

The accuracy of Eq. (S16) was reported in ref. [37] up to the first 16 eigenmodes. In Fig. 2b, we extended the analysis to show that cortical eigenmodes are highly correlated either with appropriately rotated spherical harmonics of the same order or of harmonics spanning adjacent wavelengths within the same eigengroup, up to the first 6 eigengroups comprising 49 modes, which account for ~70% of the reconstruction accuracy of fMRI task-activation maps. This correspondence explains why cortical and spherical modes perform similarly—they have very similar geometries at spatial scales relevant for reconstructing fMRI activation maps. Conversely, the idiosyncratic (non-sphere-like) attributes of cortical geometry are captured by short-wavelength, high-frequency eigenmodes.

To demonstrate specificity with respect to objects that have sufficiently distinct geometries, we obtained surface meshes for three objects from the Stanford 3D Scanning repository (http://graphics.stanford.edu/data/3Dscanrep/); an armadillo, a horse, and a dragon (Fig. 2c). We chose these objects because they have the same topology as the cortex and a sphere (i.e., they have no holes or handles), but they lack a sphere-like core and have large high-frequency perturbations (e.g., the limbs of the horse and armadillo and the spiky horns of the dragon) that represent deviations from the smooth geometric variations of the cortex and sphere. The surface meshes of the objects were regularized to ensure that they are triangular in nature with each point having approximately 6 neighbors. We then calculated the eigenmodes of these objects by solving the Helmholtz equation using the LBO, similar to how we obtained the cortical geometric eigenmodes.

We can describe the geometric similarity of the above three objects with the cortex and a sphere in two ways. In Fig. 2d, we show the normalized eigenvalue spectra of each object, which were obtained by plotting the eigenvalue of each mode normalized by the eigenvalue of the first non-zero mode (i.e., mode 2)[42]. It is evident that the eigenvalue spectra of the cortex and sphere are closely aligned, whereas the eigenvalue spectra of the other objects show sharper increases, consistent with the high-frequency fluctuations characterizing their geometries.

An alternative method to show geometric similarity is to reconstruct one object using the modes of another. Figure S4 shows the reconstructions of the surfaces of the cortex, armadillo, horse, and dragon using spherical harmonics. These reconstructions were obtained by relying on the framework of eigenmode expansion, such that

$$\begin{pmatrix} x \\ y \\ z \end{pmatrix} = \sum_{l=0}^{\infty} \sum_{m=-l}^{l} a_{lm} Y_l^m , \qquad (S17)$$

where $a_{lm}$ are the coefficient weights specific to each of the $x$-, $y$-, and $z$-coordinates and $l$ and $m$ correspond to the degree (i.e., eigengroup) and order (i.e., mode index within the eigengroup) of the spherical harmonics. Equation (S17) is similar to the one used in reconstructing empirical data, but in this case we are individually reconstructing the 3D spatial coordinates of the points on the surfaces of the objects with respect to spherical harmonics instead of reconstructing a single map. As previously shown[37], the key geometric features of the cortex can be reconstructed using just the first 49 to 100 spherical harmonics, which confirms the coarse-scale geometric similarities between the two objects and the simple linear mapping between cortical geometry and spherical modes. A much higher number of spherical harmonics is required to reconstruct the other objects, with important features still missing at 144 harmonics, underscoring their geometric dissimilarity with the sphere and cortex.

Finally, we mapped the modes of each object onto the cortical fsLR-32k CIFTI space, with 32,492 vertices in each hemisphere, via multimodal surface matching[43]. We used the mapped modes to reconstruct the 7 key HCP task-contrast maps, as was done in our original work. The results in Fig. 2e show that modes derived from the armadillo, horse, and dragon do not reconstruct fMRI activation maps as well as cortical geometric modes.

### S6. Supplementary References

1. Faskowitz, J. *et al.* Commentary on Pang et al. (2023) Nature. 2023.07.20.549785 Preprint at https://doi.org/10.1101/2023.07.20.549785 (2023).
2. Pang, J. C. *et al.* Geometric constraints on human brain function. *Nature* **618**, 566–574 (2023).
3. Wright, J. J. & Liley, D. T. J. Simulation of electrocortical waves. *Biological Cybernetics* **72**, 347–356 (1995).

4. Jirsa, V. & Haken, H. Field Theory of Electromagnetic Brain Activity. *Physical Review Letters* **77**, 960–963 (1996).
5. Robinson, P. A., Rennie, C. J. & Wright, J. J. Propagation and stability of waves of electrical activity in the cerebral cortex. *Physical Review E* **56**, 826–840 (1997).
6. Robinson, P. A., Rennie, C. J., Rowe, D. L., O'Connor, S. C. & Gordon, E. Multiscale brain modelling. *Philosophical Transactions of the Royal Society B: Biological Sciences* **360**, 1043–1050 (2005).
7. Nunez, P. L. The brain wave equation: a model for the EEG. *Mathematical Biosciences* **21**, 279–297 (1974).
8. Brodmann, K. *Physiologie des Gehirns*. (Druck der Union deutsche Verlagsgesellschaft, 1914).
9. Muller, L., Reynaud, A., Chavane, F. & Destexhe, A. The stimulus-evoked population response in visual cortex of awake monkey is a propagating wave. *Nat Commun* **5**, 3675 (2014).
10. Rubino, D., Robbins, K. A. & Hatsopoulos, N. G. Propagating waves mediate information transfer in the motor cortex. *Nat Neurosci* **9**, 1549–1557 (2006).
11. Muller, L. & Destexhe, A. Propagating waves in thalamus, cortex and the thalamocortical system: Experiments and models. *Journal of Physiology-Paris* **106**, 222–238 (2012).
12. Townsend, R. G. *et al.* Emergence of Complex Wave Patterns in Primate Cerebral Cortex. *J. Neurosci.* **35**, 4657–4662 (2015).
13. Burkitt, G. R., Silberstein, R. B., Cadusch, P. J. & Wood, A. W. Steady-state visual evoked potentials and travelling waves. *Clinical Neurophysiology* **111**, 246–258 (2000).
14. Hindriks, R., van Putten, M. J. A. M. & Deco, G. Intra-cortical propagation of EEG alpha oscillations. *NeuroImage* **103**, 444–453 (2014).
15. Muller, L., Chavane, F., Reynolds, J. & Sejnowski, T. J. Cortical travelling waves: mechanisms and computational principles. *Nat Rev Neurosci* **19**, 255–268 (2018).
16. Aquino, K. M., Schira, M. M., Robinson, P. A., Drysdale, P. M. & Breakspear, M. Hemodynamic traveling waves in human visual cortex. *PLoS Computational Biology* **8**, (2012).
17. Raut, R. V. *et al.* Global waves synchronize the brain's functional systems with fluctuating arousal. *Science Advances* **7**, (2021).
18. Robinson, P. A. Interrelating anatomical, effective, and functional brain connectivity using propagators and neural field theory. *Physical Review E* **85**, (2012).
19. Coombes, S., Beim Graben, P., Potthast, R. & Wright, J. J. *Neural fields: Theory and applications*. vol. 9783642545 (Springer, 2014).
20. Markov, N. T. *et al.* Cortical High-Density Counterstream Architectures. *Science* **342**, 1238406 (2013).
21. Henderson, J. A. & Robinson, P. A. Relations between the geometry of cortical gyrification and white-matter network architecture. *Brain Connectivity* **4**, 112–130 (2014).
22. Roberts, J. A. *et al.* The contribution of geometry to the human connectome. *NeuroImage* **124**, 379–393 (2016).
23. Wang, X. J. & Kennedy, H. Brain structure and dynamics across scales: In search of rules. *Current Opinion in Neurobiology* **37**, 92–98 (2016).
24. Robinson, P. A. *et al.* Prediction of electroencephalographic spectra from neurophysiology. *Physical Review E* **63**, 021903 (2001).
25. Rennie, C. J., Robinson, P. A. & Wright, J. J. Unified neurophysical model of EEG spectra and evoked potentials. *Biological Cybernetics* **86**, 457–471 (2002).
26. Abeysuriya, R. G., Rennie, C. J. & Robinson, P. A. Physiologically based arousal state estimation and dynamics. *Journal of Neuroscience Methods* **253**, 55–69 (2015).

27. Robinson, P. A. Integrals and series related to propagators of neural and haemodynamic waves. *Royal Society Open Science* **8**, 211562 (2021).
28. Coombes, S., Lord, G. J. & Owen, M. R. Waves and bumps in neuronal networks with axo-dendritic synaptic interactions. *Physica D: Nonlinear Phenomena* **178**, 219–241 (2003).
29. Deco, G., Jirsa, V. K., Robinson, P. A., Breakspear, M. & Friston, K. The dynamic brain: From spiking neurons to neural masses and cortical fields. *PLoS Computational Biology* **4**, (2008).
30. Braitenberg, V. & Schüz, A. *Cortex: Statistics and Geometry of Neuronal Connectivity*. (Springer-Verlag Berlin, 1998).
31. Robinson, P. A. Physical brain connectomics. *Phys. Rev. E* **99**, 012421 (2019).
32. Markov, N. T. *et al.* Weight Consistency Specifies Regularities of Macaque Cortical Networks. *Cerebral Cortex* **21**, 1254–1272 (2011).
33. Robinson, P. A. Neural field theory of effects of brain modifications and lesions on functional connectivity: Acute effects, short-term homeostasis, and long-term plasticity. *Phys. Rev. E* **99**, 042407 (2019).
34. Alexander-Bloch, A. F. *et al.* On testing for spatial correspondence between maps of human brain structure and function. *NeuroImage* **178**, 540–551 (2018).
35. van Essen, D. C. *et al.* The WU-Minn Human Connectome Project: An overview. *NeuroImage* **80**, 62–79 (2013).
36. Barch, D. M. *et al.* Function in the human connectome: Task-fMRI and individual differences in behavior. *NeuroImage* **80**, 169–189 (2013).
37. Gabay, N. C. & Robinson, P. A. Cortical geometry as a determinant of brain activity eigenmodes: Neural field analysis. *Physical Review E* **96**, (2017).
38. Robinson, P. A. *et al.* Eigenmodes of brain activity: Neural field theory predictions and comparison with experiment. *NeuroImage* **142**, 79–98 (2016).
39. Wagner, H. H. & Dray, S. Generating spatially constrained null models for irregularly spaced data using Moran spectral randomization methods. *Methods in Ecology and Evolution* **6**, 1169–1178 (2015).
40. Burt, J. B., Helmer, M., Shinn, M., Anticevic, A. & Murray, J. D. Generative modeling of brain maps with spatial autocorrelation. *NeuroImage* **220**, 117038 (2020).
41. Vos de Wael, R. *et al.* BrainSpace: a toolbox for the analysis of macroscale gradients in neuroimaging and connectomics datasets. *Commun Biol* **3**, 1–10 (2020).
42. Reuter, M., Wolter, F. E. & Peinecke, N. Laplace-Beltrami spectra as 'Shape-DNA' of surfaces and solids. *CAD Computer Aided Design* **38**, 342–366 (2006).
43. Robinson, E. C. *et al.* Multimodal surface matching with higher-order smoothness constraints. *NeuroImage* **167**, 453–465 (2018).

### S7. Supplementary Figures

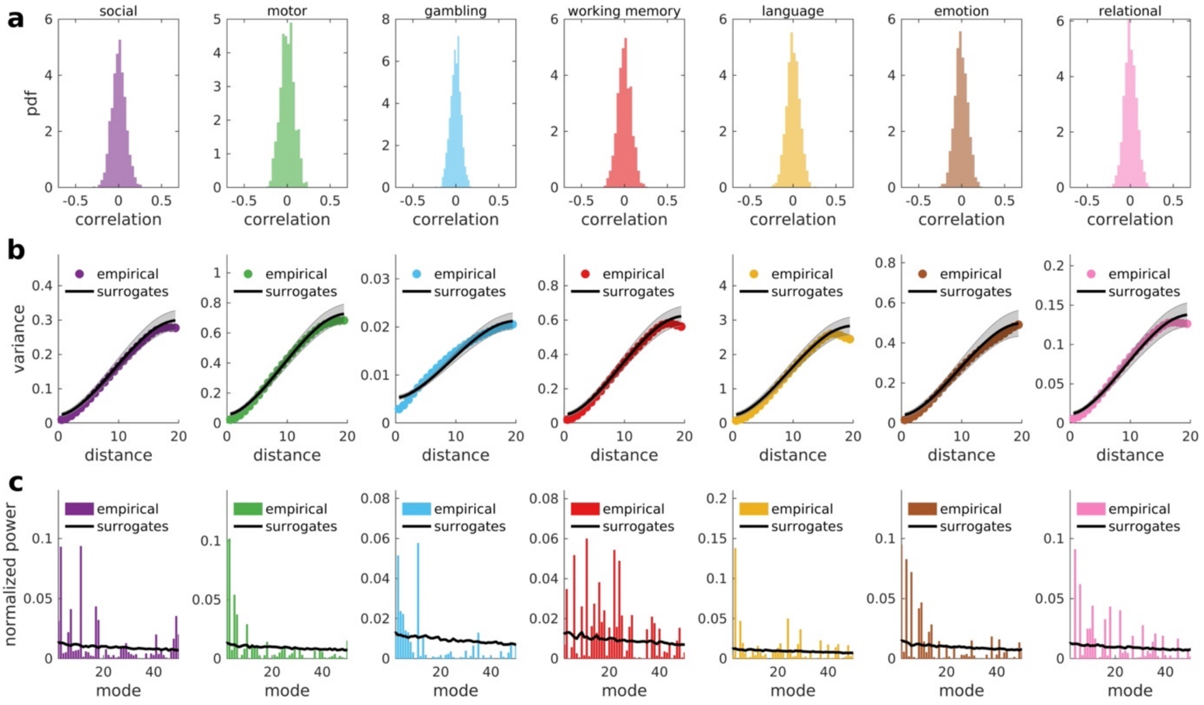

**Figure S1. Statistics of SA-preserving parametric null surrogates of the 7 key HCP task-contrast maps. (a)** Distributions of correlations between the empirical map and 1000 surrogate maps. The distributions are centered at zero with narrow tails (compared to the much broader distributions produced by the Moran spectral randomization, as shown in Fig. S4 of Faskowitz et al.[1] and Supplementary Fig. 7 in ref. [40]), indicating that the parametric surrogates properly randomize the empirical maps. (**b**) Variograms (variance vs distance) of empirical and surrogate maps. The surrogates preserve autocorrelations of empirical data at spatial scales relevant for capturing low-level fMRI processing effects. (**c**) Normalized power spectra of empirical and surrogate maps. The surrogates appropriately destroy the spatial wavelength structure of empirical data.

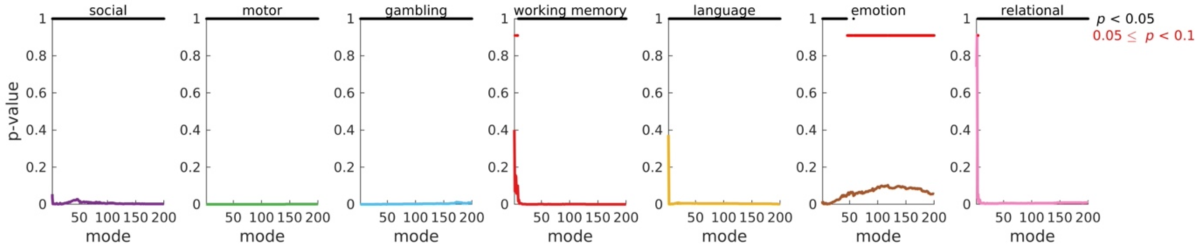

**Figure S2. Statistical test for reconstructing the 7 key HCP task-contrast maps.** The panels show the $p$-values, representing the proportion of times the accuracy of geometric eigenmodes in reconstructing empirical data exceeds the accuracy in reconstructing 1000 surrogate maps for each mode. The black asterisks indicate $p < 0.05$ (uncorrected for multiple comparisons) and the red asterisks indicate $0.05 < p < 0.1$. The results clearly show the effectiveness of geometric eigenmodes against null data.

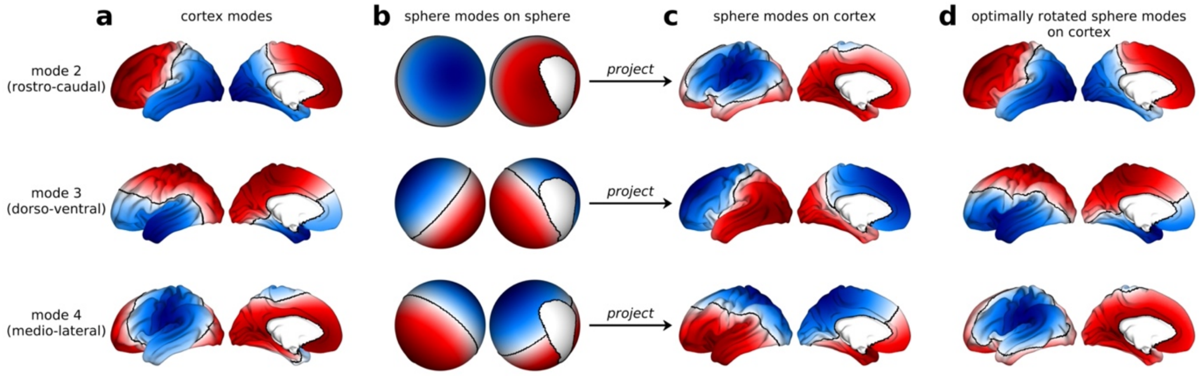

**Figure S3. Relationship between cortex and spherical modes of the first eigengroup. (a)** Cortical modes for the first eigengroup have a single node (black line) fixed by the cortical geometry. Hence, these modes represent specific spatial variations along the rostro-caudal, dorso-ventral, and medio-lateral directions. (**b**) Spherical modes on the spherical surface have arbitrary orientations because of their degenerate (i.e., equivalent) eigenvalues, meaning that the modes have no preferred orientation—they are rotationally invariant (Fig. 1b). (**c**) Spherical modes naively projected on the cortical surface without any alignment to the cortical modes. The nodal lines do not match the real ones in panel **a** imposed by the cortical geometry. (**d**) Spherical modes projected on the cortical surface after rotation to maximally align them with the cortical modes. Due to the degeneracy of the spherical modes, each one can be rotated so that it aligns with any of the cortical modes. This example thus shows the equivalence between coarse-scale cortical and spherical modes when appropriate rotations are applied.

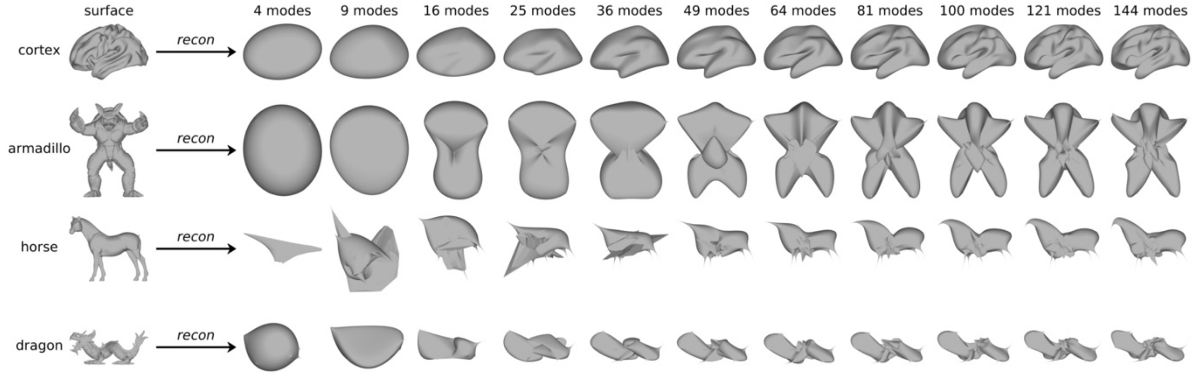

**Figure S4. Reconstruction of cortical and non-cortical surface geometries using spherical modes.** The x, y, and z spatial locations of points on the surface are individually reconstructed in terms of increasing number of spherical modes [Eq. (S17)]. The coarse-scale shape of the cortex is already captured by about 49 to 100 spherical modes, with more intricate characteristics such as the sulci and gyri becoming more well-defined as more modes are used. This result further demonstrates the geometrical similarity of the cortex and a sphere. The other objects cannot be easily reconstructed by spherical modes, showing that their geometrical features are very dissimilar to a sphere and cortex, which highlights the suitability of their eigenmodes as benchmarks for the cortical geometric eigenmodes.
